# Spatial and temporal alterations in protein structure by EGF regulate cryptic cysteine oxidation

**DOI:** 10.1101/624304

**Authors:** Jessica B Behring, Sjoerd van der Post, Arshag D Mooradian, Matthew J Egan, Maxwell I Zimmerman, Jenna L. Clements, Gregory R Bowman, Jason M Held

**Author notes:** These authors contributed equally to this work (Jessica B Behring, Sjoerd van der Post, Arshag D Mooradian). Corresponding Author: Jason M. Held, Washington University School of Medicine, Campus Box 8076, 660 South Euclid Avenue, Saint Louis, MO 63110. PNAS Classification: Biological Sciences – Biochemistry.

## Abstract

Stimulation of receptor tyrosine kinases (RTK) such as EGF locally increase reactive oxygen species (ROS) levels at the plasma membrane that oxidize cysteines in proteins to enhance downstream signaling. Spatial confinement of ROS is an important regulatory mechanism to redox signaling, but it remains unknown why stimulation of different receptor tyrosine kinases (RTKs) at the plasma membrane target distinct sets of downstream proteins. To uncover additional mechanisms specifying which cysteines are redox regulated by EGF stimulation, we performed time-resolved quantification of the oxidation of 4,200 cysteine sites subsequent to EGF stimulation in A431 cells. EGF induces three distinct spatiotemporal patterns of cysteine oxidation in functionally organized protein networks, consistent with the spatial confinement model. Unexpectedly, protein crystal structure analysis and molecular dynamic simulation indicate widespread redox regulation of cryptic cysteines that are only solvent exposed upon changes in protein conformation. Phosphorylation and increased flux of nucleotide substrates serve as two distinct modes by which EGF specifies which cryptic cysteines become solvent exposed and redox regulated. Since proteins structurally regulated by different RTKs or cellular perturbations are largely unique, solvent exposure and redox regulation of cryptic cysteines is an important mechanism contextually delineating redox signaling networks.

**Significance Statement:** Cellular redox processes are interconnected, but are not in equilibrium. Thus, understanding the redox biology of cells requires a systems-level, rather than reductionist, approach. Factors specifying which cysteines are redox regulated by a stimulus remain poorly characterized but are critical to understanding the fundamental properties of redox signaling networks. Here, we show that EGF stimulation induces oxidation of specific cysteines in 3 distinct spatiotemporal patterns. Redox regulated proteins include many proteins in the EGF pathway as well as many cysteines with known functional importance. Many redox regulated cysteines are cryptic and solvent exposed by changes in protein structure that were induced by EGF treatment. The novel finding that cryptic cysteines are redox regulated has important implications for how redox signaling networks are specified and regulated to minimize crosstalk. In addition, this time-resolved dataset of the redox kinetics of 4,200 cysteine sites is an important resource for others and is an important technological achievement towards systems-level understanding of cellular redox biology.

## Introduction

Activation of many cell surface receptors transiently increase reactive oxygen species (ROS), predominantly hydrogen peroxide (H_2_O_2_), that act as important signaling second messengers. Growth factor activation of receptor tyrosine kinases (RTKs) are the best studied examples and include EGF (3), insulin (24), PDGF (84), IGF-1 (30), FGF (77), and NGF (97), but cytokines (53, 56, 87), B- and T-cell receptors (46, 71), integrins (85), and GPCRs (27) are cell surface receptors that also increase ROS production upon activation. ROS-dependent cellular phenotypes are pleiotropic including cell migration, proliferation, differentiation, polarization, and cell death (8).

Oxidation of cysteines is a key mechanism by which ROS transduce signaling changes. Oxidative inhibition of the catalytic cysteine of protein tyrosine phosphatases (24, 33, 47) and in the case of EGF, redox regulation of EGFR itself (65), functionally contribute to signaling. The factors that specify which cysteines are oxidized by ROS produced in response to a stimulus are therefore the critical determinants regulating the specificity and crosstalk of redox signaling networks.

Spatial restriction of ROS (15,50,54,65,92,93) within subcellular microdomains (15) is an important contributor determining which proteins and cysteines are oxidized. In the case of EGF, the best studied redox signaling pathway, this occurs through localized activation of NADPH oxidases and inactivation of peroxiredoxins (PRDXs) at the plasma membrane (15, 50, 54, 65,81,92). As solvent accessibility and pKa of a cysteine are key determinants of its oxidizability, in the canonical model EGF stimulation locally oxidizes solvent accessible cysteines with low pKa within endosomal microdomains (13, 15, 50, 65,92, 93). Notably, the molecular details of cysteine oxidation upon EGF stimulation are static (65, 96) and it remains unknown how spatiotemporally dynamic cysteine redox is upon EGF stimulation. Elucidating the dynamic downstream redox control of proteins on a global scale at different points during the course of EGFR and NADPH Oxidase (NOX) internalization and trafficking therefore requires a new approach.

Despite the clear importance of spatial regulation to redox signaling, it is not the only factor determining which cysteines are oxidized by a stimulus (10, 14, 64). For example, while EGF and insulin both generate NOX-derived ROS at the plasma membrane via activation of the RTKs, EGFR and the insulin receptor, respectively, PTP1b redox regulation is highly sensitive to insulin but poorly oxidized by EGF (33, 65). This differential redox sensitivity can be extended broadly as unique patterns of PTPs are redox regulated upon stimulation by different RTKs and other cell surface receptors (10, 57, 58, 64). Since these are all spatially constrained similarly at the cell surface, it is unclear what additional factors specify which cysteines are oxidized by specific stimulus or context and how redox signaling pathways at the cell surface are buffered from one another.

We therefore performed a workflow termed OxRAC for cysteine <u>ox</u>idation analysis by <u>r</u>esin-assisted capture that advances existing redox proteomic workflows (28, 29) to characterize the dynamics of cysteine redox networks by coupling enrichment of oxidized cysteines with high resolution, data-independent acquisition mass spectrometry analysis (DIA-MS) for peptide quantitation. We quantify time-resolved changes in the oxidation state of 3,566 unique cysteine-containing peptides covering 4,200 cysteine sites at five timepoints after EGF stimulation in A431 cells, a common cellular model for EGF signaling studies (65, 96). We identified three cysteine redox networks with distinct spatiotemporal regulation and functional organization. Notably, protein structure analysis and molecular dynamics simulation reveals that cryptic cysteines (2), those that are only solvent exposed upon changes in protein conformation, are unexpectedly widespread and important contributors governing the specificity of redox signaling networks. EGFR-dependent phosphorylation and increased nucleotide substrate flux serve as two distinct mechanisms by which EGF specifies which cryptic cysteines are redox regulated.

## Results

### The OxRAC workflow to globally profile dynamic changes in cysteine oxidation

Serum starved A431 cells were treated with 100 ng/mL EGF 0, 2, 5, 15, 30 and 60 minutes prior to lysis in a degassed lysis buffer with NEM and SDS to fully denature proteins and block free thiols (Fig. 1A). This differential alkylation strategy is a common way to preserve the redox state of cysteines (29, 35) and facilitates time-resolved kinetic analysis. It also limits alterations in cell signaling that occur when cysteine sulfenic acids are labeled *in situ* in cells as previously performed to investigate EGF-dependent cysteine oxidation (65, 96). A 1hr incubation with the SOH targeting probe Dyn-2 (65), typical for EGF redox studies (96), alters EGF-dependent signaling on its own, increasing phosphorylation of ERK 3.9 fold and global phosphotyrosine 2.1 fold (Supp. Fig S1). Rapid alkylation by NEM and inclusion of EDTA in the lysis buffer minimizes non-specific oxidation (29, 32, 35), as demonstrated in this study by the low median percent oxidation of cytoplasmic proteins (10.2%) which is consistent with previous reports (31). In addition, complete protein unfolding and alkylation by NEM was verified by Western blot (Supp. Fig. S2 A). For OxRAC analysis, samples are then reduced and proteins with previously oxidized thiols are bound to thiopropyl sepharose resin (28, 29). Spike in of an isotope-encoded, cysteine-containing peptide was performed prior to binding to normalize for differences in resin binding across samples (Fig. 1B). Proteins are digested on-resin and peptides containing the previously oxidized cysteines are retained on the resin, eluted by reduction, alkylated with iodoacetamide (IAC), and analyzed by LC-MS using both data-dependent acquisition (DDA) to identify peptides and DIA-MS for quantitation using high resolution MS2 chromatograms (Fig. 1B) (72). OxRAC enriches and analyzes only the oxidized cysteines to increase coverage depth and incorporates DIA-MS analysis for high precision quantitation with limited to no missing data (12). This solves a limitation of isobaric tagging approaches for cysteine redox proteomics, which often have 25% or more data missing (11, 79) and thus have limited statistical power especially when multiple hypothesis correction appropriate for large proteomics datasets is considered (29, 49, 79, 96).

**Figure 1.**
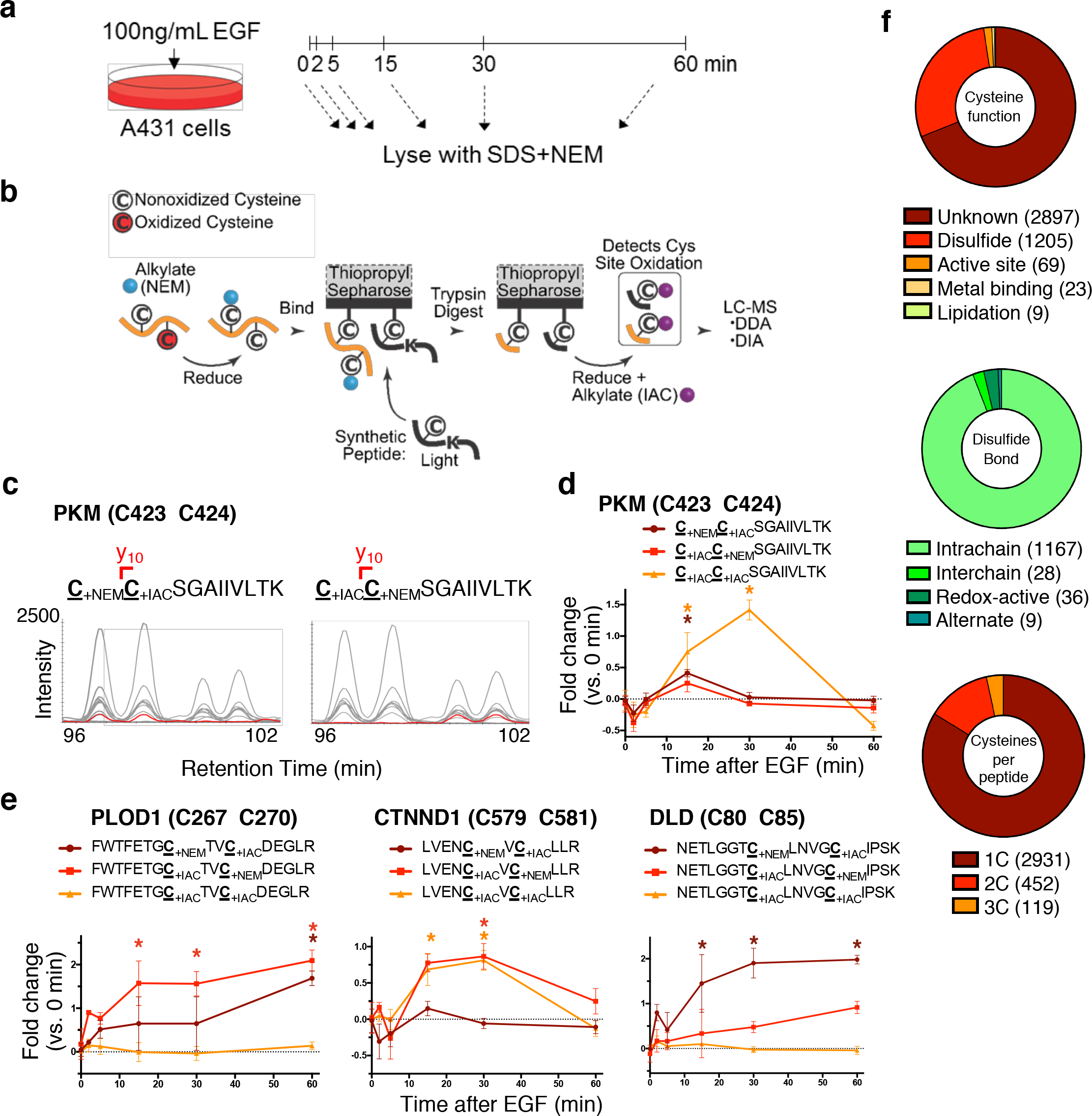
The OxRAC workflow to globally profile cysteine oxidation and overview of results. **A)** Serum-starved A431 cells were left untreated (0 min) or stimulated with 100ng/mL EGF for the times indicated prior to lysis. **B)** OxRAC workflow schematic in which free cysteines are trapped with NEM and oxidized thiols are enriched by thiopropyl sepharose resin and trypsin digested on-resin. The oxidized cysteines remain bound during washing, then are eluted by reduction and labeled with iodoacetamide (IAC) to differentiate oxidized (IAC-labeled) from non-oxidized (NEM-labeled) cysteines. Peptides are analyzed by data-dependent acquisition (DDA) to identify peptides and data-independent acquisition (DIA) mass spectrometry to quantify based on high resolution MS2 scans. **C)** DIA-MS2 scans of the pyruvate kinase (PKM) peptide CCSGAIIVLTK. The site defining y10 fragment ion (red line) between the two labeled cysteines confirms peak identity. **D)** Time-dependent changes in the relative oxidation of PKM C423 and C424, and cysteines in **E)**procollagen-lysine 2-oxoglutarate 5-dioxygenase 1 (PLOD1), catenin delta-1 (CTNND1), and dihydrolipoyl dehydrogenase (DLD) in response to EGF stimulation. **F)** Enumeration of the functional annotation, disulfide bond types, and number of cysteines per peptide in the dataset. Asterisks indicate significant changes at specific time points based on ANOVA (Dunnet’s post-hoc test) and error bars are SEM.

Two control samples were analyzed in parallel to confirm minimal false peptide identifications, remove background signal, and estimate the stoichiometry of oxidation of each cysteine. Samples that were fully reduced and alkylated (‘RED-NEM’) prior to digestion, resin binding, and IAC treatment served as a negative control. RED-NEM samples identified very few IAC modified peptides (<0.05%) which confirms minimal false peptide identifications (28), and exhibited minimal signal intensity by LC-MS (Supp. Fig S2 B). This signal was subtracted from each peptide to remove background and improve quantitative accuracy. Samples reduced prior to thiopropyl sepharose (‘RED’) account for protein expression and are used to estimate the oxidation stoichiometry of each cysteine.

DIA often differentiates which cysteine is oxidized in a peptide containing multiple cysteines (Fig. 1C). The extracted fragment ion chromatogram for the pyruvate kinase (PKM) peptide CCSGAIIVLTK with two adjacent cysteines demonstrates how this method distinguishes peptide pairs (Fig. 1C). These cysteines can be individually quantified using the *m/z* of the y10 fragment ion (red, Fig. 1C) as well as label-specific changes in relative hydrophobicity and retention times; NEM introduces a chiral center that resolves as peak doublets in reversed phase chromatography (25). Each singly oxidized PKM cysteine exhibits a similar temporal response to EGF, yet only C424 was significantly redox regulated (Fig. 1D). The doubly oxidized C267, C270 peptide was also significantly oxidized, maximally at 30 minutes after EGF stimulation (Fig. 1E). Additional examples of differentially regulated peptides with cysteines in close proximity are found in PLOD1, CTNND1, and DLD (Fig. 1E).

The redox status was quantified for 4,200 cysteine sites in 3,566 cysteine-containing peptides. 99% of cysteines were unambiguously assigned to a specific cysteine residue including 94.5% of peptides containing multiple cysteines. The functional annotation of almost 70% of the identified sites is unknown (Fig. 1F). Of those annotated, 1205 sites participate in disulfide bonding, the majority of which are intramolecular (Fig. 1F).

### EGF-dependent regulation of cysteine redox networks cluster into three distinct temporal profiles associated with unique subcellular locations and biological processes

To discern if there was time-dependent regulation of cysteine redox networks, the distribution of fold-changes versus baseline at time 0 for all cysteine-containing peptides was examined (Fig. 2A). Widespread EGF-dependent oxidation was observed at 15 and 30 minutes as indicated by right shifted, non-Gaussian distributions. To median center and normalize these samples, Gaussian mixture modeling was used to best fit normal distributions (Fig. 2A, red lines) to the data. They were then centered according to the median of the left-most distribution in each sample since EGF increases ROS production and cysteine oxidation (3, 22, 65).

**Figure 2.**
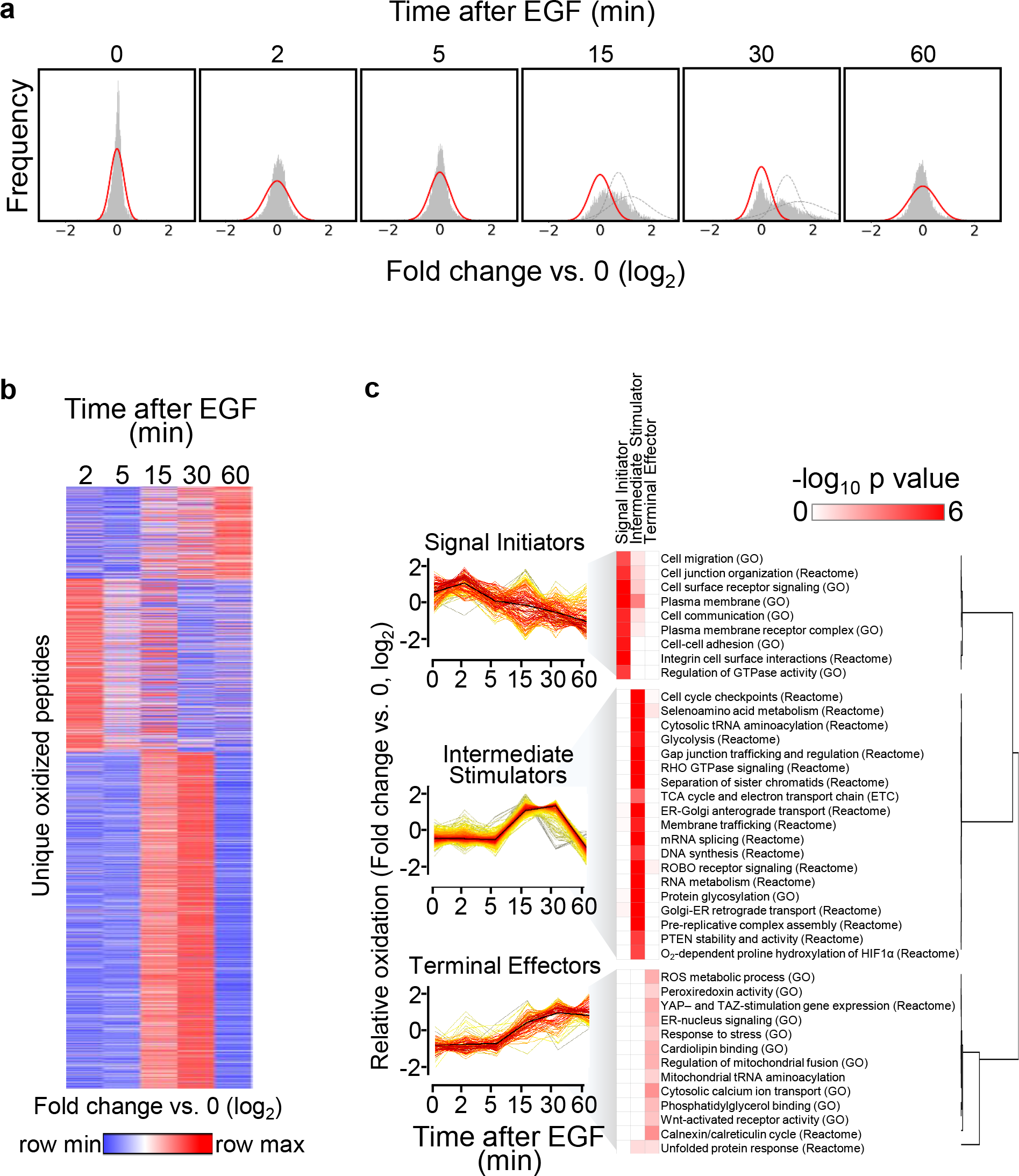
EGF-dependent regulation of cysteine redox networks cluster into three distinct temporal profiles associated with unique subcellular locations and biological processes. **A)** Log_2_ fold change of all peptides versus baseline (average of three time 0 replicates). Lines indicate normal distributions and the red line indicates the fold change used for normalization. **B)** Heatmap of all cysteine-containing peptides clustered by K-means (1 minus pearson correlation, K = 3) of relative oxidation levels. **C)** Fuzzy c-means clustering of significantly oxidized peptides and selected Gene Ontology (GO) and Reactome annotations.

Strikingly, over half of the cysteines quantified (51.3%) were significantly oxidized by EGF (ANOVA, Benjamini-Hochberg corrected, q<0.05). While high, the magnitude of oxidation of the cysteine redoxome is consistent with other published endogenous perturbations including an increase in sulfenation of 49% percent of cysteine sites in A431 cells upon EGF stimulation (96), 52% percent of cysteines oxidized by light-dark cycles (29), and 60% of cysteines oxidized upon fasting (49).

To characterize the kinetics of EGFR-dependent redox regulation of protein networks we performed 3-component K-means clustering (Fig. 2B). Three distinct groups were observed without over-partitioning and most cysteines were maximally oxidized at 15-30 minutes, consistent with their non-normal fold-change distributions (Fig. 2A). The remaining peptides clustered either by peaking immediately at 2 minutes or having a delayed increase. We performed fuzzy c-means clustering to further define the temporal pattern of cysteine oxidation. Partitioning the data into 5 clusters indicated three unique temporal patterns of redox regulation (Fig. 2C), which we term “signal initiators”, “intermediate stimulators”, and “terminal effectors” using previously coined time-based descriptors of EGFR regulation (63). The “signal initiators” and “terminal effectors” appeared as separate clusters whereas “intermediate stimulators” encompassed the remaining 3 clusters, which we combined into a single representative cluster.

The biological roles of each of the three cluster were inferred by performing Gene Ontology (GO) enrichment analysis. Consistent with the prevailing model of localized ROS production by EGF, cysteines were regulated spatiotemporally, in concert with EGFR trafficking upon EGF glycosylation, nucleotide processing as well as phosphatase and HIF biology. “Terminal effectors” were associated with the cell periphery (Fig. 2C, Supplementary 24 Table S2). “Intermediate stimulators” were enriched for metabolism, trafficking, protein were involved in stress response, ROS metabolism, peroxiredoxin activity, calcium transport, and ER-nucleus signaling, which may indicate negative feedback. Surprisingly, EGF stimulation redox regulated many aspects of mitochondrial biology; TCA and electron transport chain, mitochondrial fusion, as well as cardiolipin binding. Taken together, EGF regulates cysteine redox networks with three distinct temporal profiles that are associated with a wide range of biological processes.

### Cysteines in all major organelles are oxidized by EGF but location influences the temporal dynamics

Prompted by the time-resolved regulation of spatially distinct subcellular compartments, we more systematically investigated the spatial regulation of EGF-dependent cysteine oxidation. Cytoplasmic and extracellular or luminal cysteines exhibited different temporal profiles of oxidation. Cytoplasmic cysteines peaked at 30 minutes and were less oxidized at 60 minutes whereas oxidation of extracellular and luminal localized cysteines was sustained over 60 minutes (Fig. 3A). Two cysteines in the cell surface receptor PLXNB2, C1484 and C937, localized to the cytoplasmic and extracellular sides of the cell, respectively, showed distinct patterns of redox regulation wherein the extracellular cysteine was rapidly and transiently oxidized, similar to “signal initiators”, and its intracellular cysteine was maximally oxidized at 15 and 30 minutes and back to baseline at 60 minutes, in parallel with “intermediate stimulators” (Fig. 3B).

**Figure 3.**
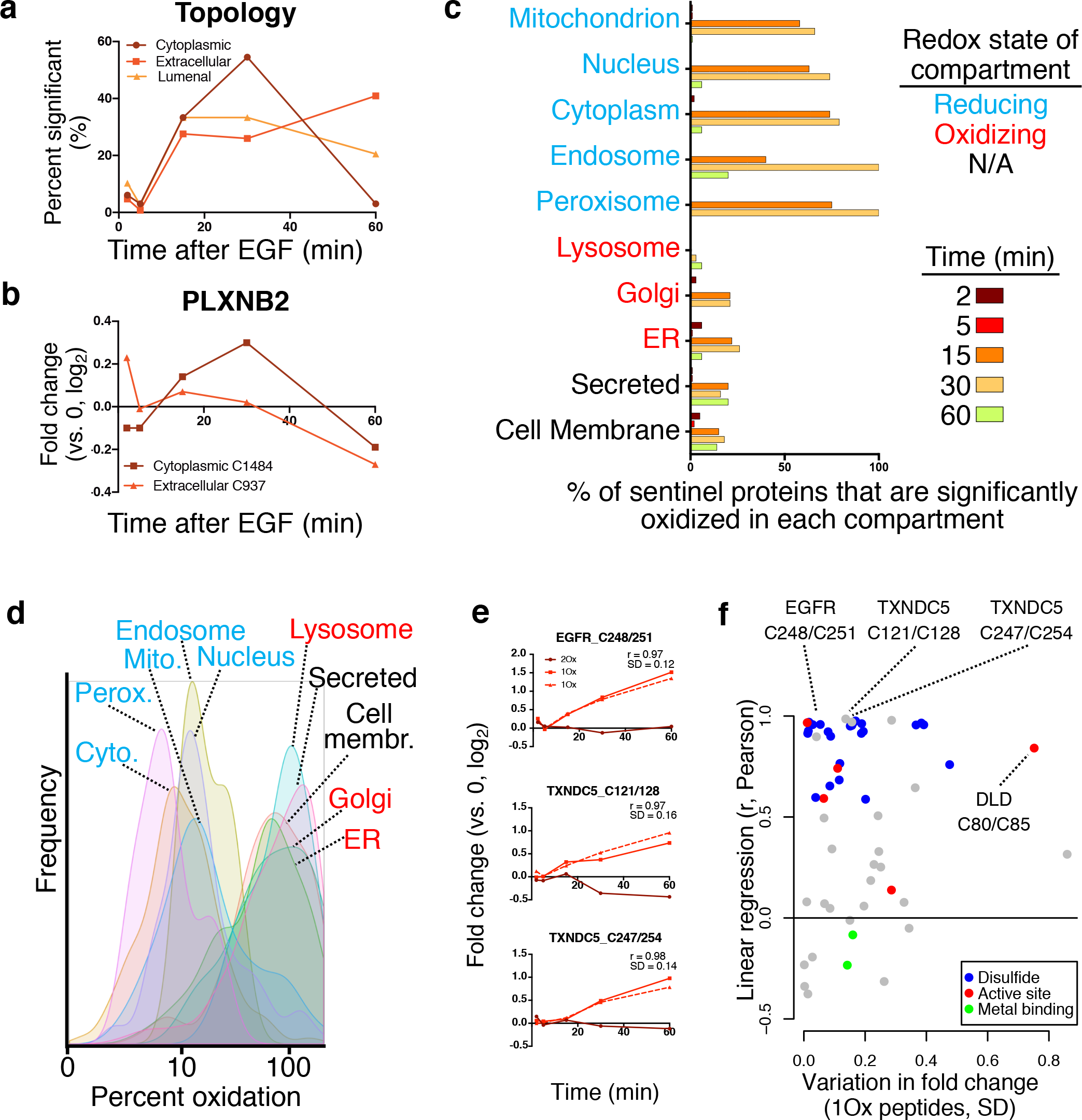
Cysteines in all major organelles are oxidized by EGF but location influences the temporal dynamics. **A)** Membrane orientation of 182 modified sites on either the cytoplasmic, extracellular, or lumenal side presented as percentage significant over time in response to EGF. Significant per time point is based on p<0.01 by ANOVA with Benjamini Hochberg post-hoc test correction. **B)** Differential response over time of the extracellular and cytoplasmic side of the transmembrane protein Plexin-2B. **C)** The percentage of cysteines in 942 sentinel proteins detected by OxRAC and annotated to a single cellular compartment that have at least one cysteine significantly oxidized by EGF. **D)** Estimated percent oxidation of all cysteines in sentinel proteins. **E)** Selected examples of differentially oxidized peptides containing 2 cysteines. A known disulfide bond in EGFR between cysteine 248 and 251 becomes reduced upon endocytosis in response to EGF. Examples of two unknown functional cysteines in TXNDC5 which are potentially disulfide linked. 1Ox and 2Ox indicate singly or doubly oxidized forms of the peptide, respectively. **F)** Linear regression (r, Pearson) of the fold-change over time of the singly oxidized (1Ox) forms versus the variation (standard deviation, SD) between the two singly oxidized sites. Sites annotated as disulfide linked, active, or metal binding are indicated. Includes differentially oxidized sites identified in peptides spanning two cysteines.

To add organelle precision to characterization of redox localization, we assessed changes in cysteine oxidation of 943 “sentinel” proteins detected that were annotated in Uniprot to only a single subcellular compartment (Supp. Table S1). EGF stimulation increased the percent of sentinel proteins oxidized in each of 10 cellular organelles, further establishing that EGF dependent redox regulation extends beyond plasma membrane microdomains (Fig. 3C). Sentinel proteins at the cell membrane were redox regulated most rapidly (Fig. 3C), consistent with GO annotations of “signal initiators”.

Sentinel proteins in organelles that are predominantly reducing had a higher likelihood of being oxidized in response to EGF than those in oxidizing compartments (Fig. 3C). To determine if this was due to higher basal oxidation of cysteines in oxidizing compartments, limiting the potential for increased oxidation, we estimated the percent oxidation for peptides at baseline using samples prepared in parallel that were reduced prior to thiopropyl-sepharose to quantify total peptide levels in the sample. As expected, proteins in oxidized compartments were highly oxidized even at steady state, in contrast to reduced compartments (Fig. 3D). Notably, peroxisomes are major cellular sources of ROS and often considered an oxidizing environment, however, our analysis indicates sentinel proteins in peroxisomes are both highly responsive to EGF-dependent ROS and have a basal oxidation percentage similar to reducing compartments (Fig. 3D). These results demonstrate that EGF redox regulates cysteines in all major subcellular compartments but the magnitude of oxidation increase is limited by the basal redox potential of each compartment.

Temporal patterns of cysteine redox regulation can distinguish specific redox processes and cysteine oxoforms. Extracellular cysteines are predominantly disulfide linked, and upon EGF-induced internalization these cysteines are reduced in endosomes (95). Most peptides annotated as ‘disulfide’ showed a unique temporal pattern as shown in Figure 3E. Levels of singly oxidized peptides (1Ox) increased linearly together over the course of 60 minutes, along with a small decrease or no change in the level of the doubly oxidized (2Ox) peptide, consistent with internalization and partial reduction in endosomes throughout the time course. This is a unique pattern, distinguishable from those annotated as active sites such as DLD C80/C85 in which only one cysteine was specifically increased (Fig. 1E). These two regulatory hallmarks were observed for several non-annotated cysteines that group with known disulfide-linked cysteines, including TXNDC5 C121/128 and C247/254 and others (Fig. 3F), and suggests that these are novel disulfide-linked cysteines.

### Synchronized redox regulation of cysteines throughout canonical EGF signaling pathways at 15 and 30 minutes

Redox regulation of cysteines is prevalent in many members of canonical EGF-driven signaling pathways including matrix metalloproteinases, Rho family GTPases, and proteins involved in adherens junctions, protein translation, and proliferation (Fig. 4, Supp. Table S3). Components of caveolar-mediated endocytosis, which controls plasma membrane recycling and degradation of EGFR itself, are also redox regulated. Redox regulation was largely synchronous across most EGF-related pathways, peaking at 15-30 minutes after EGF stimulation as “intermediate stimulators”. Notably, caveolar-mediated endocytosis and matrix metalloproteinases were two pathways with a greater proportion of reduced cysteines, each of which include proteins that are endocytosed into the reducing environment of endosomes.

**Figure 4.**
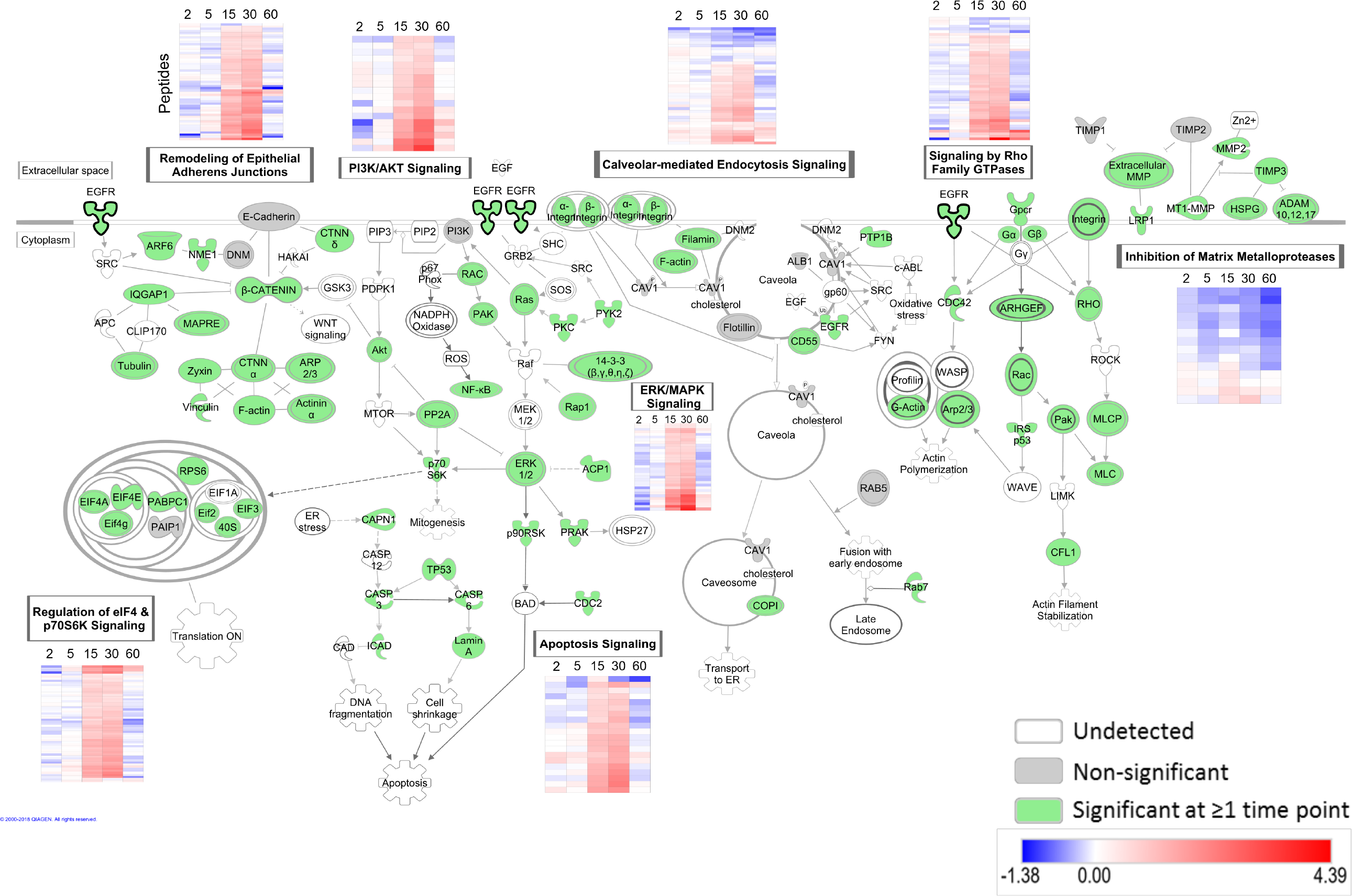
Synchronized redox regulation of cysteines throughout canonical EGF signaling pathways at 15 and 30 minutes. Select enriched canonical pathways from IPA downstream of EGFR are pictured. All genes with a significantly changing cysteine are colored green. Proteins detected but not significantly oxidized by EGF are filled in grey and those undetected, but important for continuity of a pathway were left unfilled. EGF redox dynamics over 60 minutes for significantly changing peptides in each pathway are represented in heatmaps.

Oxidation of tyrosine phosphatases and kinases is hypothesized to enhance EGFR-dependent phosphotyrosine signaling which peaks rapidly in A431 cells at ~1 minute (22, 65). However, the 31 unique peptides that are assigned to protein phosphatases and 44 unique peptide sequences assigned to kinases primarily have much slower temporal kinetics of “intermediate stimulators” and maximally redox regulated at 15 and 30 minutes (Supp. Fig. S3). This includes catalytic cysteines in the well-known redox regulated protein protein-tyrosine phosphatase 1B (PTP1B) (47, 65) and ACP1 (also known as LMW-PTP), which is capable of dephosphorylating phosphotyrosines in proteins (100). While ACP1 oxidation by EGF has not been previously reported and may play an underappreciated role in modulating EGF-dependent, phosphorylation the substantial delay of redox regulation of phosphotyrosine modifying enzymes raises questions about its mechanistic role in EGFR-dependent phosphotyrosine signaling.

### Protein domains redox regulated by EGF stimulation

While canonical redox active protein domains were detected by OxRAC, many cysteines in these domains were not significantly redox regulated following EGF stimulation (Fig. 5A). The C-terminal domain of 1-cys peroxiredoxins (“1-cysPrx_C”), containing the resolving cysteine, were not redox regulated in response to EGF (Fig. 5A). Closer inspection of proteins containing the 1-cysPrx_C domain revealed that none of the resolving cysteines in PRDXs 1-4 were oxidized by EGF (Fig. 5B). While the AhpC-TSA domain containing both the peroxidatic and allosteric regulator cysteines appeared to be EGF-regulated (Fig. 5A), this was solely due to significant changes in oxidation of allosteric sites rather than the peroxidatic cysteine (Fig. 5B). Peroxidatic cysteines in PRDX5 and 6 were quantified, but neither were significantly oxidized (Fig. 5B, lower panel). However, unlike the canonically redox active cysteines, allosteric regulator cysteines in PRDX1-4 exhibited up to a nearly 16-fold change in oxidation (Fig. 5B).

**Figure 5.**
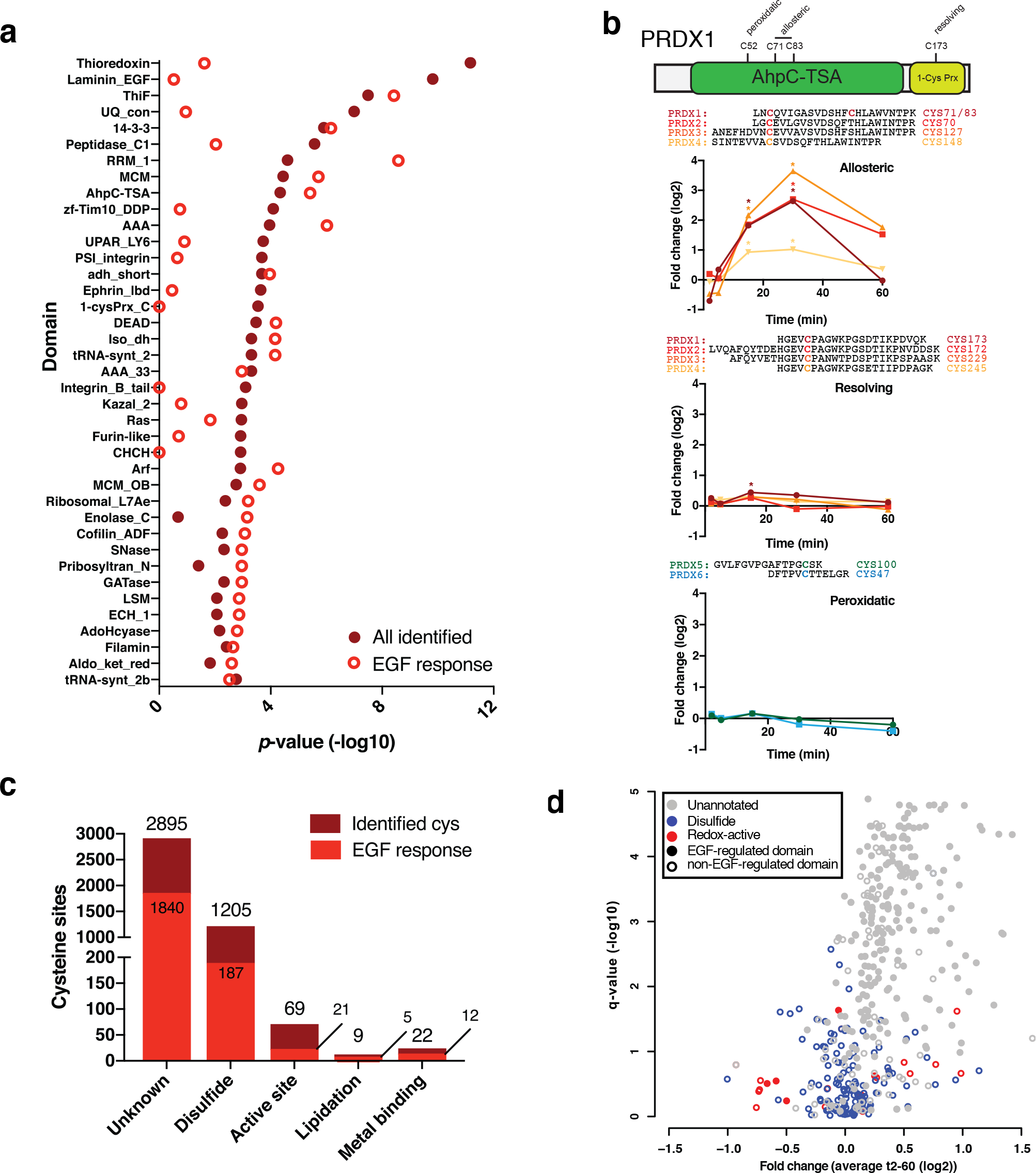
Protein domains redox regulated by EGF stimulation. **A)** Enrichment of protein domains detected in the entire dataset ‘all identified’ or significantly oxidized by EGF, ‘EGF response’, as compared to all domains in the human proteome. P-values determined by two-tailed Fisher’s exact test. **B)** Domain organization of PRDX1 and its cysteine locations and function. The relative oxidation of the allosteric, resolving and peroxidatic cysteines in PRDXs over time in response to EGF stimulation. Asterisks indicate significant changes at specific time points based on ANOVA (Dunnet’s post-hoc test). **C)** Functional annotation of all cysteines detected compared to those significantly regulated in response to EGF. **D)** The average ratio of all peptides assigned to the most enriched domains vs. their significance (q), overlaid with functional annotation when available. Closed circles: enriched sites in response to EGF, open circles: not enriched in response to EGF.

Other protein domains and families were hotspots for EGF-dependent redox regulation (Fig. 5A) including the RNA recognition motif (RRM_1), AAA ATPases, 14-3-3 proteins, small GTPases, and kinases. One example of pervasive regulation throughout a protein family is AAA ATPases, comprising 57 genes with very low cysteine sequence conservation that play a diverse role in cellular functions (Supp. Fig S4 A). OxRAC profiling detected that 33 of 37 peptides assigned to ATPase family members were significantly oxidized by EGF (Supp. Fig. S4 A, Fisher’s *p*=10e^−6^). One of these AAA ATPases, Valosin-containing protein (VCP), is the most oxidized protein in this study, with EGF-redox regulated cysteines throughout all of its domains (Supp. Fig. S4 B), including VCP C522 which inhibits VCP function (61). Since VCP plays a key role in resolving ER stress, and EGFR signaling is enhanced by ER stress (99), oxidative inactivation of VCP upon EGF stimulation may enhance EGFR signaling.

### Functional analysis of the redox regulated cysteines

The EGFR signaling network is robust, parallelized, and pleiotropic, thus even canonical downstream regulators such as AKT and mTOR are dispensable or limited contributors to EGF-induced phenotypes (19, 36, 89). Complex feedback loops compensate for loss of a single regulatory effector (18, 74) complicating the reductionist approach of site-directed mutagenesis which also not applicable to enzymes in which the cysteine residue is directly involved in their activity (42, 43). To our knowledge no single point mutation of a post-translational modification site of canonical downstream effectors substantially effects EGF-induced phenotypes.

Therefore, to delineate if EGF-dependent redox regulation has a functional impact on protein function we first examined if active sites or other functionally relevant cysteines were significantly oxidized by EGF treatment. These include the phosphatases ACP1 and PTP1B which require their redox regulated active sites C13 (86) and C215 (98), respectively, caspases-1 and −3 which are inactivated by mutation of C215 (60) and C163 (6) respectively, GAPDH which requires C152 (73), and OTUB1 which is catalytically inert when C91 is mutated (41). P4HB is unable to bind a misfolded substrate with mutation of the significantly redox regulated C397 and C400 (76), cathepsin B is inactivated by mutation of the significantly redox regulated C108 (48), UGDH has negligible activity when the significantly redox regulated C276 is mutated (17), VCP C522 is required for activity (61), and thioredoxin which is unable to release substrates when the significantly redox regulated C32 and C35 are mutated (52). In addition, the yeast homolog of IDI1 is much less catalytically active upon mutation of its significantly redox regulated active site C86 (82).

We next investigated if cysteines oxidized by EGF stimulation are allosteric regulators of protein function. AKT C296 is significantly oxidized by EGFR treatment and is an allosteric inhibitory site in AKT (91). Mutation of C183 in ERK, which is significantly oxidized, abrogates its phosphorylation and activity as well as diminishes ERK’s antiapoptotic potential in response to nitrosative stress (21). Together, these findings verify that many cysteines oxidized by EGFR are functionally relevant. Notably, only 11% of the cysteines significantly oxidized following EGF stimulation are annotated in Uniprot as an active site, disulfide bond, metal binding, or lipidated (Fig. 5C), indicating that many of the identified sites may have allosteric functions (Fig. 5D, filled grey circles).

### Redox-independent regulation of protein structure by EGF specifies cryptic cysteines for oxidation

Both solvent accessibility and pKa are potential determinants of a cysteine’s likelihood to be oxidized. Bioinformatic prediction using the surrounding amino acid sequence and available structural information (68) revealed that cysteines significantly redox regulated by EGF have no change in pKa (Supp. Fig. S5). Furthermore, most of the oxidized cysteines have a low relative solvent accessibility (RSA), even slightly lower than those that were detected but not oxidized (Fig. 6 A, B). The median RSA of cysteines oxidized by EGF is 6% (Fig. 6A), well below the threshold of 25% that is considered solvent accessible (68). This surprising observation is consistent with Yang *et al.* (96) since evaluating their dataset with the same solvent prediction tool found that the cysteine targets of exogenous H_2_O_2_ were largely solvent accessible, as expected, but those targeted by EGF were largely solvent inaccessible (96) (Fig. 6C). The unexpected lack of solvent accessibility of EGF-dependent redox regulated cysteines led us to further investigate what governed the oxidation of these solvent inaccessible cysteines.

**Figure 6.**
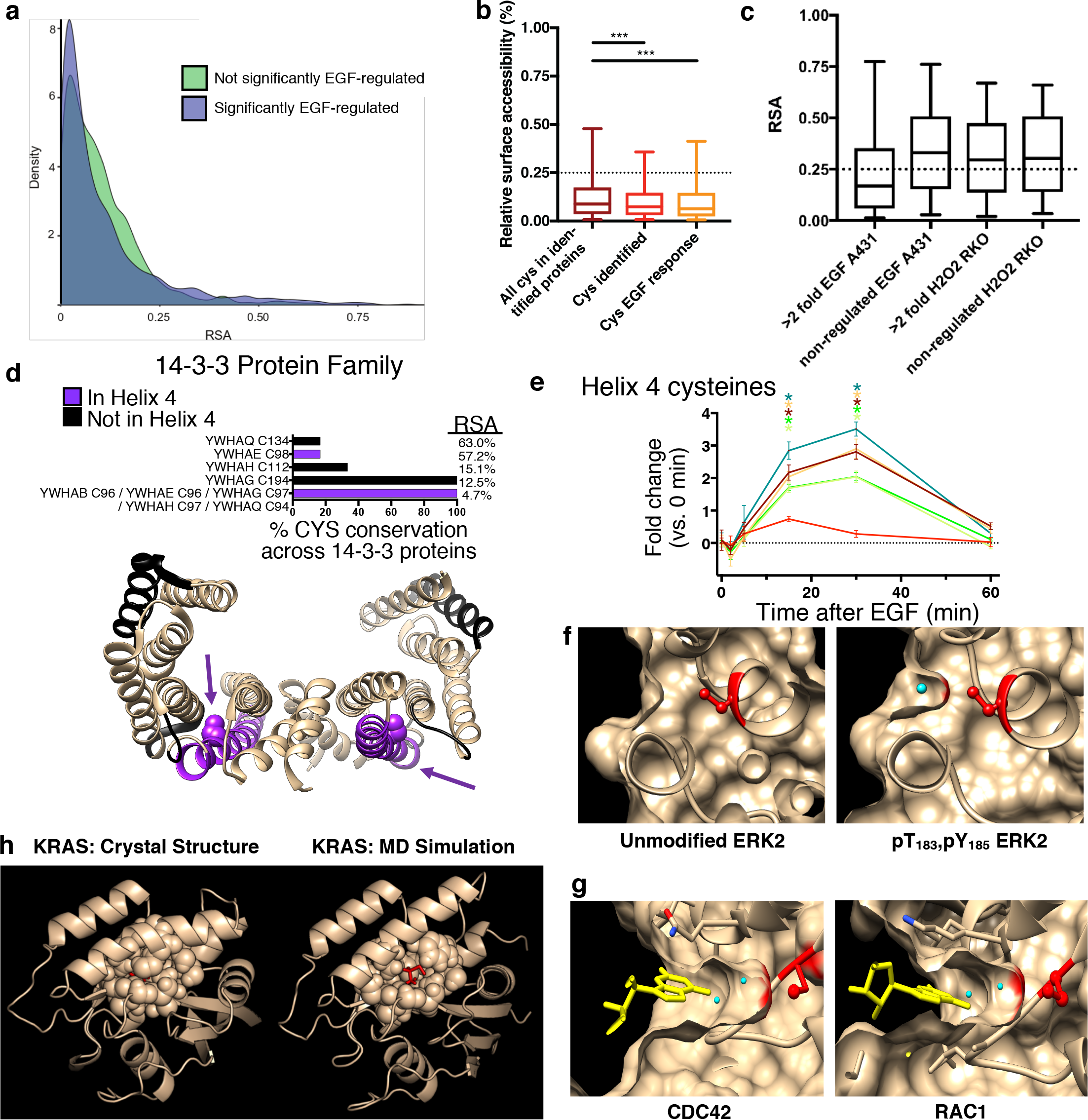
EGF-dependent redox regulation of buried cysteines in 14-3-3, small GTPase proteins and ERK2. **A)** Density plot of RSA comparing cysteines significantly and not regulated by EGF. **B)** Relative solvent accessibility (RSA) for all cysteines in proteins identified compared to all cysteines specifically quantified and those significantly regulated. Sites >0.25 considered solvent accessible. * p< 0.05, *** p <0.001 by one-way ANOVA. For the box-whisker are functional annotation when available. Closed circles: enriched sites in response to EGF, open plot: center line is median, limits are upper and lower quartiles, and whiskers are 5 and 95 percentiles. **C)** RSA prediction for sulfenated cysteines in (96). **D)** Amino acid sequence conservation of the cysteines identified in 14-3-3 proteins. Structure of dimerized 14-3-3 sigma (PDB: 4DAU) with helix 4 shown in purple and locations of the remaining oxidized cysteines shown in black. Purple arrows point to the conserved cysteine in Helix 4, represented as a sphere. Relative surface accessibility (RSA) is indicated for each cysteine site or the average across 14-3-3 family members. **E)**Time-dependent changes in the oxidation of 14-3-3 cysteines. Asterisks indicate significant changes at specific time points based on ANOVA (Dunnet’s post-hoc test) and error bars are SEM. **F)** Unmodified and phosphorylated (T183 and Y185) ERK2 crystal structures (PDB: 1ERK and 2ERK, respectively) with Cysteine 65 highlighted in red and solvent as a cyan sphere. **G)** Crystal structures of the GTP binding pocket in CDC42 (PDBID: 5CJP GTP analogue, peptide YVEC[+57]SALTQK) and Rac1 (PDBID: 5O33 GTP analogue, peptide YLEC[+57]SALTQR). C157 is highlighted in red. Yellow structure is the bound GTP nucleotide analogue. Cyan spheres indicate solvent molecules. **H)** Representative crystal structure of KRAS (left, PDB: 5VQ2) in which C80 (red) is solvent inaccessible as well as a representative structure from MD simulations of KRAS (right) in which C80 is solvent accessible. Amino acid side chains within 7 angstroms of C80 are shown as spheres, and C80 is shown in red as a ball-and-stick

The 14-3-3 protein family consists of 7 highly conserved genes that bind phospho-serine/phospho-threonine motifs, interact with EGFR, and promote EGFR signaling (62, 67). 8 of 9 peptides detected in 14-3-3 proteins were significantly oxidized by EGF (Fisher’s *p*=7e^−7^), most notably oxidation of a highly conserved cysteine in helix 4 in five of the six 14-3-3 family members detected (Fig. 6 D, E). This helix 4 cysteine is one of the least solvent accessible amino acids in 14-3-3 proteins, as determined from crystal structures, and substitution to either alanine or serine destabilizes the protein (39). Consistent redox regulation of this buried cysteine across nearly all 14-3-3 proteins verifies that highly solvent inaccessible cryptic cysteines (7) are *bona fide* hotspots for EGF-dependent redox regulation.

We therefore hypothesized that redox-independent regulation of protein conformation as a consequence of EGF stimulation may solvent expose cryptic cysteines for oxidization and provide context-dependence. One potential mechanism by which EGF may induce conformational changes is protein phosphorylation. ERK C65 is significantly oxidized by EGF (~1.96-fold at 30 min, q=0.0016), but is solvent inaccessible in the unmodified ERK crystal structure with an RSA of 4.2% (Fig. 6F). However, the crystal structure of ERK2 phosphorylated at T183 and Y185, a consequence of EGF stimulation (66, 75), shows that C65 becomes solvent exposed (Fig. 6F, blue sphere indicates solvent). Mutation of this cysteine has been shown to confer a level of resistance to ERK inhibitors, with one study suggesting that site may be close enough in proximity to affect the ATP binding pocket which these drugs target (23).

Many members of the Small GTPases family play a role in EGFR signaling. OxRAC profiling identified 20 peptides, assigned to 18 genes, in the small GTPase family that were significantly oxidized by EGF (Fisher’s *p*=0.01). Many diverse small GTPase family members were significantly oxidized, though the closely related RHO, RAC, and CDC42 proteins, known to play a role in EGFR-dependent endocytosis, invasion, and metastasis, were preferentially targeted (Supp. Fig. S4 C). These results indicate that the small GTPase family is far more redox-sensitive than previously described (59). Examination of crystal structures for CDC42 and Rac1 show that the thiol of the cryptic cysteines oxidized by EGF (C157 in each protein, RSAs of 2.5 and 2.6%, respectively) its solvent accessible at the base of the pocket is blocked by the nucleotide ligand (Fig. 6 G, yellow sticks). As EGF increases the GTPase activity of these proteins (45), increased nucleotide exchange would enable H_2_O_2_ to oxidize the cysteine. Therefore EGF-dependent changes in enzyme activity is an additional mechanism by which EGF-dependent stimulation specifies which cryptic cysteines are redox regulated.

Analysis of VCP crystal structures further validate EGF-dependent oxidation of cryptic cysteines. VCP C77, which is significantly oxidized (Supp. Fig. 4 B) is solvent accessible in the ATP?S bound structure but solvent inaccessible in the ADP bound state (Supp. Figure 4C). Another VCP cysteine, C209, is significantly oxidized by EGF stimulation and lies at the base of the nucleotide ligand pocket similar to GTPases and would be solvent accessible during turnover. Taken together, cryptic cysteines in multiple proteins are oxidized by EGF via structural changes induced by phosphorylation, activity, and nucleotide flux.

There are only limited examples of phosphoprotein crystal structures, and crystal structures are biased towards low energy conformers and don’t accurately convey all the structural ensembles of a protein (80). KRAS is activated by EGF and transduces signaling to ERK, but EGF only increased the oxidation of solvent inaccessible KRAS C80 rather than solvent accessible C118 which is known to be redox regulated (51). To find evidence of cryptic pockets in KRAS that expose cryptic cysteines, we characterized its protein conformational ensembles with the FAST algorithm to observe transitions between states that are inaccessible to conventional molecular dynamics (MD) (102, 103). C80 has an exposed solvent accessible surface area (69) in approximately 4% of the states, which arises from a cryptic pocket opening between helices 2 and 3 (representative conformation shown in Fig. 6H). This transient solvent exposure of a cryptic, buried cysteine provides a potential mechanism for redox reactions to occur, supporting the plausibility of the widespread redox regulation of cryptic cysteines that our experimental data suggest.

## Discussion

The EGF pathway is an intensively characterized and modeled signaling network due to its importance in migration, differentiation, and cancer (44,63,78,88,94,96). However, our current understanding of the dynamics and specificity of cysteine redox regulation is limited compared to extensively characterized phosphorylation, acetylation, ubiquitination and interactome changes stimulated by EGF treatment of A431 cells (44, 63, 88, 94,101). We therefore we quantified redox regulation of the cysteine redoxome at a systems level with OxRAC to address three major questions: (i) is redox regulation by EGF stimulation confined to endosomal microdomains; (ii) what specifies which cysteines are oxidized by EGF, but not other stimuli with similar spatial redox regulation such as other growth factors; (iii) what is the temporal correlation and interplay between downstream cysteine oxidation and phosphorylation?

A total of 3,566 unique cysteine-containing peptides spanning 4,200 cysteine sites were quantified by OxRAC for three biological replicates at each of six timepoints. The high depth of coverage was evidenced by the detection of low-abundant proteins such as p53, which previously required targeted MS analysis for detection (35), and included many functionally relevant cysteines. EGF stimulation significantly oxidized more than half (51.3%) of the peptides in our dataset, a percentage on par with increased sulfenation of 49% of cysteines upon EGF stimulation (96) and other physiological stimuli that oxidize nearly half of cysteines detected (29, 49).

EGF stimulation induces clear spatiotemporal regulation of the cysteine redox networks. Three temporal patterns of cysteine redox regulation were identified, each with unique biological and functional organization (Fig. 2). Unexpectedly, these were located throughout all major cellular organelles (Figs. 2 & 3) demonstrating that EGF-dependent redox signaling is not confined to endosomal microdomains. Several possibilities can explain the unexpectedly broad spatial distribution of oxidized cysteines. One is that ROS diffuse further into the cell than previously thought (15, 92) and that localized microdomains of ROS—such as those in the nucleus that emerge via phosphorylation-mediated inhibition of PRDX1 near the centrosome (50)—occur throughout the cell to facilitate cysteine oxidation. Second, redox active endosomes may translocate very broadly throughout the cell. Third, direct oxidation of cysteines by ROS may occur proximal to redox active endosomes, but disulfide-relays, EGF-dependent changes in the redox potential of an organelle, or translocation of a protein from one redox environment to another expand the reach of cysteine oxidation throughout the cell.

The kinetics of the cysteine redoxome after EGF stimulation challenges the canonical regulatory model in which redox regulation of tyrosine phosphatases and kinases is an important amplifier of phosphotyrosine signaling. Cysteines in phosphotyrosine modifying enzymes are primarily oxidized 15 and 30 minutes after EGF stimulation, matching the observed kinetics of peak ROS production and sulfenation upon EGF stimulation (22, 65)), but far delayed from phospho-tyrosine signaling which peaks at ~1 minute post-EGF stimulation in multiple studies (13, 63). Given this temporal delay, it remains unclear how redox regulation of phosphotyrosine modifying enzymes affects EGF-dependent phosphorylation. To our knowledge there no kinetic analyses have established that redox inactivation of PTPs is synchronized with the rapid increase in phospho-tyrosine.

The substantial delay between phosphotyrosine signaling and redox regulation, does support an emerging regulatory model of redox signaling in which redox-independent changes in protein structure and activity, those resulting from specific stimuli such as EGF, specify which cysteines are oxidized via changes in solvent accessibility (2). Our results demonstrate that EGF oxidizes many cryptic cysteines that are not solvent accessible in crystal structures (Fig. 6a), thus this regulation is widespread. This observation is consistent with Gould et al. who observed that cysteines oxidized at steady state in mouse liver are buried (26) and Yang et al. (96), who found that while the sulfenated targets of H_2_O_2_ were largely solvent accessible, but the cysteine targets sulfenated by EGF have a median surface accessibility of only 16%—well below the accessible cutoff of 25% (Fig. 6C,(68)). Protein crystal structures suggest at least two possible mechanisms by which EGF stimulation can regulate the solvent accessibility of redox regulated cryptic cysteines: phosphorylation and GTP/GDP cycling. Regulatory domains of PTPs have previously been shown to contribute to oxidation sensitivity PTPs (64) including SHP2 whose cryptic catalytic cysteine is blocked by an SH2 domain. When a RTK becomes phosphorylated, SHP-2’s SH2 domain extends to bind the receptor which subsequently allows the cryptic catalytic cysteine to become oxidized (5, 58). Together, these results establish that EGF-dependent, redox-independent conformational regulation of cryptic allosteric cysteines is a widespread mechanism that confers context-dependent selectivity in redox signaling networks.

These data suggest a novel role for allosteric rather than the peroxidatic and resolving PRDX cysteines in EGFR signaling (Fig. 5B) that builds on others findings that catalytic and resolving cysteines of PRDXs are substantially less sensitive to EGF treatment than H_2_O_2_ (13, 47, 93, 96). There is no increased oxidation of the peroxidatic cysteine in PRDX5 and PRDX6 (Fig. 5B), suggesting that these sites do not play a role in disulfide relays (81, 90) upon EGF treatment. Despite the lack of oxidation of the canonical residues in PRDXs, we observe that allosteric sites in PRDXs are highly redox regulated (Fig. 5B). To our knowledge, no study has demonstrated functional dependency of PRDX disulfide relays on the peroxidatic cysteine. We therefore suggest that PRDX allosteric sites may be key mediators of disulfide relays upon EGF stimulation, consistent with evidence that these allosteric PRDX cysteines form disulfide linked heterodimers with other proteins (38).

The goal of redox systems biology is to characterize the dynamics of redox signaling pathways in an integrative and comprehensive manner (34). The singularly unique connections of the cysteine redoxome to cellular redox networks, coupled with the ability to assess or infer their position in protein structure, subcellular location, and function, can be leveraged to provide an integrative portrait of cellular redox regulation (34). OxRAC analysis and cysteine redox proteomics can profile the system-wide dynamics of redox regulation in time and space, integrate information about redox regulation mediated directly by and independent of ROS, and uniquely profile context-dependent redox regulation due to changes in protein conformation and activity. This work provides a database of potentially druggable cysteines (4), including many cryptic cysteines unexpected to be oxidized by EGF, and shifts the paradigm of what governs specificity in redox signaling networks. In addition, as many proteins canonically downstream of EGFR are redox regulated (Figs. 4–6), these results offer new avenues for integrating redox signaling into current therapeutic interventions.

## Materials & Methods

### Material

All materials are from Sigma unless otherwise noted.

### Cell culture and treatments

A431 cells were maintained in DMEM with 4.5 g/L glucose+1mM pyruvate and 10% FBS (Gibco). 2e6 cells were plated in 10cm dishes, grown for 3 days, then serum starved overnight. Zero time points as well as samples that would be fully reduced or fully alkylated as positive and negative controls, respectively, were not treated with EGF. For all others, Human recombinant EGF protein (R&D Systems) was diluted to 10x in serum free medium, added to each dish (final concentration of 100 ng/mL) and swirled. After the appropriate incubation time (2, 5, 15, 30, or 60 min), media was aspirated and lysis buffer added.

### Differential thiol alkylation

Cells were lysed in 500 *μ*L degassed, fully denaturing lysis buffer (3% SDS, 10mM EDTA, 200mM Tris pH 7.0, 100 mM NEM except for the Reduced and Reduced+NEM samples that were lysed without NEM but in the presence of 11 mM TCEP to reduce). Proteins in Reduced samples are unblocked and therefore all can bind to thiopropyl sepharose, whereas cysteines in Reduced + NEM lysates are full alkylated (Supp. Fig S2) and are only bound and detected due to non-specific binding. Lysates were probe sonicated briefly then incubated at 37 C for 45 min at 950 rpm to alkylate. For the RED+NEM samples, lysates were reduced at 50 C for 45 min followed by addition of 100 mM NEM to the for 30 min at 37 C for 45 min. All samples were quenched after NEM incubation with 133 mM cysteine (60 min, 37 C, 950 rpm). All samples were then reduced with 11 mM TCEP (45 min, 50 C, 950 rpm) and precipitated with 4 volumes of methanol overnight at −80 C.

### In situ dyn-2 treatment and Western blots

A431 cells were grown and serum starved as for EGF treatment and OxRAC prior to addition of 5 mM dyn-2 (Acme Bioscience) or vehicle (1% DMSO) for 1 hr. Cells were lysed in 3% SDS, 25 mM Tris pH 7.4 and assayed by Western blot. Antibodies were from Cell Signaling except for pan-phosphotyrosine (pY) which was from Millipore. Densitometry was performed with Image J.

### Preparing thiopropyl sepharose 6B

20 mg Thiopropyl sepharose 6B resin (GE Life Science) per sample was washed with 1 mL degassed water for 15 min to rehydrate at room temperature. The supernatant was removed and 1 mL water was added to wash resin. Resin was centrifuged (1000 × g, 30 sec) and washed again with 1 mL water. This process was repeated twice more, but washed with thiopropyl sepharose binding buffer (6 M guanidine HCl, 0.1% NP-40, 25 mM Tris pH 7.4, 150 mM NaCl, 5 mM EDTA). Buffer was aspirated prior to addition of protein lysates.

### Thiopropyl sepharose 6B binding

Precipitated proteins were centrifuged (1,500 × g, 5 min) and the supernatant was removed. Pellets were washed with 1 mL 80% MeOH three times. Samples were resuspended in 800 *μ*L thiopropyl sepharose binding buffer (6 M guanidine HCl, 0.1% NP-40, 25 mM Tris pH 7.4, 150 mM NaCl, 5mM EDTA) using sonication and vortexing. 4 pmol unlabeled CysOx Peptide Standard (VSAPGAGSAKADTGPACGTAR, custom synthesized by ThermoFisher) was added to each sample for normalization. This synthetic peptide has a non-human sequence and contains a cysteine to bind to the resin and an internal lysine that is cleaved by trypsin and released enabling normalization during sample processing. Lysates were added to thiopropyl sepharose resin and incubated in the dark at 4 C rotating end-over-end.

### Thiopropyl sepharose 6B washing and elution

All buffers were degassed prior to use and all washes are 500uL, rotating end over end, for 10 min at room temperature. Lysates plus resin were transferred to spin columns (Sigma) and centrifuged to remove unbound protein. The resin was washed three times with thiopropyl sepharose binding buffer, three times with QCET buffer (50 mM Tris pH 8.0 plus 1 mM EDTA), 4 times with SDS/NP-40 buffer (1% NP-40, 0.1% SDS, 5mM EDTA, 150 mM NaCl, 25 mM Tris pH 7.4), 4 times with 80% acetonitrile, and 4 times with high salt buffer (2 M NaCl, 50 mM Tris pH 7.4). to prepare for trypsin digestion, resin was briefly washed with trypsin digestion buffer (1 M urea, 50 mM Tris pH 8.0). 4 *μ*g sequencing grade trypsin (Roche) was dissolved in 165 *μ*L ice cold trypsin digestion buffer and added to resin in the spin columns with the bottom capped. Digestion proceeded at 37 C, 950 rpm overnight. Spin columns were centrifuged to elute non-bound peptides, which contain non-oxidized peptides in proteins with an oxidized cysteine, and can be analyzed separately. Resin was washed three times with 6 M guanidine HCl, 5 mM EDTA, 150 mM NaCl, 25 mM Tris pH 7.4, five times with high salt buffer, and 3 times with 80% acetonitrile, and 3 times with QCET buffer.

Peptides were eluted with 100 *μ*L 20 mM DTT in QCET buffer by capping the bottom of the spin column and incubating lysate at 60 C, 900 rpm, for 10 min. Samples were briefly centrifuged to collect eluent, and elution was repeated 2 additional times for 10 min each. 100 *μ*L 80% acetonitrile was used as a final elution. 150 *μ*L of 100 mM iodoacetamide in QCET buffer was added to eluents and incubated in the dark for 45 min at room temperature and was quenched by addition of 3 *μ*L 1 M cysteine in QCET buffer for 30 min at room temperature. Formic acid was added to acidify samples for oasis sample cleanup. Samples were vacuum centrifuged to near dryness, acidified with 0.5% formic acid and desalted using HLB Oasis SPE cartridges. Eluted samples were concentrated to near dryness and resuspended in 0.5% formic acid for LC-MS.

### LC-MS

Samples were analyzed by reverse-phase HPLC-ESI-MS/MS using a nano-LC 2D HPLC system (Eksigent) which was directly connected to a quadrupole time-of-flight (QqTOF) TripleTOF 5600 mass spectrometer (AB SCIEX) in direct injection mode. 3 *μ*L of analyte was loaded onto 3 *μ*l sample loop. After injection, peptide mixtures were transferred onto a self packed (ReproSil-Pur C18-AQ, 3*μ*m,Dr. Maisch) nanocapillary HPLC column (75 *μ*m I.D. × 22 cm column) and eluted at a flow rate of 250 nL/min using the following gradient: 2% solvent B in A (from 0-7 min), 2-5% solvent B in A (from 7.1 min), 5-30% solvent B in A (from 7.1-130 min), 30-80% solvent B in A (from 130-145 min), isocratic 80% solvent B in A (from 145-149 min) and 80-2% solvent B in A (from 149-150 min), with a total runtime of 180 min including mobile phase equilibration. Solvents were prepared as follows: mobile phase A: 0.1% formic acid (v/v) in water, and mobile phase B: 0.1% formic acid (v/v) in acetonitrile. Mass spectra were recorded in positive-ion mode. After acquisition of ~1-3 samples, TOF MS spectra and TOF MS/MS spectra were automatically calibrated during dynamic LC-MS & MS/MS autocalibration acquisitions injecting beta-galactosidase (AB SCIEX). Automatic re-calibration of TOF-MS and TOF-MS/MS scans (after every 1-3 samples) guaranteed reliably high mass accuracy and MS detector precision over an extended period of time (weeks) without interruption of the HPLC-MS/MS acquisitions. Two different mass spectrometric acquisition workflows were performed in this study: **1) Data dependent acquisitions (DDA)**: for collision induced dissociation tandem mass spectrometry (CID-MS/MS), the mass window for precursor ion selection of the quadrupole mass analyzer was set to ± 0.7 *m/z*. MS1 scans ranged from 380-1250 *m/z* at a resolution of 30,000 with an accumulation time of 250 ms. The precursor ions were fragmented in a collision cell using nitrogen as the collision gas. Advanced information dependent acquisition was used for MS2 (MS/MS) collection on the TripleTOF 5600 at a resolution of 15,000 to obtain MS2 spectra for the 50 most abundant parent ions following each survey MS1 scan. Dynamic exclusion features were based on value M not *m/z* and were set to an exclusion mass width 50 mDa and an exclusion duration of 30 sec. MS2 scans ranged from 100-1500 *m/z* with an accumulation time of 50 msec. **2) Data independent MS2 acquisitions (DIA)**: In the ‘SWATH’ DIA MS2 acquisition, instead of the Q1 quadrupole transmitting a narrow mass range through to the collision cell, a wider window of ∼10 m/z is passed in incremental steps over the full mass range (*m/z* 400-1250 with 85 SWATH segments, 63 msec accumulation time each, yielding a cycle time of 5.5 sec which includes one MS1 scan with 50 msec accumulation time). SWATH MS2 produces complex MS/MS spectra that are a composite of all the analytes within each selected Q1 *m/z* window.

### Bioinformatic database searches for TripleTOF 5600

Mass spectral data sets were analyzed and searched with both MaxQuant (http://www.biochem.mpg.de/5111795/maxquant) and Mascot (Matrix Science) against the Uniprot Human Reference Proteome. Search parameters MaxQuant included: First peptide search tolerance of 0.07 Da and main peptide search tolerance of 0.0006 Da, and variable methionine oxidation, protein N-terminal acetylation, carbamidomethyl, and NEM modifications with a maximum of 5 modifications per peptide. 2 missed cleavages and trypsin/P protease specificity. Razor protein FDR was utilized and the maximum expectation value for accepting individual peptides was 0.01 (1% FDR) and a minimum score for modified peptides of 25. For all Mascot searches, parameters were the same except for mass tolerance of 25 ppm and 0.1 Da for MS1 and MS2 spectra, respectively, and decoy searches were performed choosing the *Decoy* checkbox within the search engine. For all further data processing, peptide expectation values were filtered to keep the FDR rate at 1%.

### Skyline Data Analysis

Skyline software (https://skyline.ms/project/home/begin.view?) was used to manually examine and quantify DIA data. Spectral libraries were generated in Skyline using the DDA database searches of the raw data files. Raw files were directly imported into Skyline in their native file format and only cysteine containing peptides were quantified.

### Data submission to public database

The RAW and processed data associated with this manuscript will be deposited to the ProteomeXchange repository with the identifier PXD010880.

### Data processing

Each run was normalized for loading by dividing all raw intensities by the respective run’s light peptide raw intensity. Swath runs for each sample were averaged then every peptide modified sequence was background-subtracted by subtracting the respective peptide’s bio-rep average for RED+NEM. Any data points below background were considered to be zero. In the case of multiple charge states for a single peptide modified sequence, values were summed. Values from peptides with missed cleavages were also summed if the missed cleavage peptide did not span any additional cysteines and matched to the same exact gene(s) as the fully cleaved peptide. To calculate fold-change, the derived values from each of 3 individual biological replicates at 2, 5, 15, 30, and 60 min were divided by the average of the zero time point bio-reps. These were log_2_ transformed and then carried forward for further analysis.

### Estimating Percent oxidation of individual peptides

Following loading normalization and background subtraction, sample signal intensities for each oxidized peptide were divided by the Reduced signal intensity, which estimates the total amount of each peptide (free and reversibly oxidized forms) in these cells.

### Fitting normal distributions for re-normalization of relative expression changes

Gaussian mixture models were made using the GaussianMixture function from the scikit-learn package (v.0.19.1) in Python (v.3.6). Diagonal covariance matrices were used to constrain the models, and a maximum of 100 iterations were used for fitting. For timepoints at 2, 5, and 60 min of EGF treatment, the data from each replicate were modeled and centered using one component. For the 15 and 30 min timepoints, three components were used to identify Gaussian subpopulations, and the modeled peak with the least fold change was used for centering the data. Data were plotted using matplotlib (v.2.2.3).

### Statistical analysis

1-way ANOVA was calculated for each unique peptide modified sequence (three bio-reps for six time points; F-statistic, df 5, 12). Two-sided Dunnett’s post-hoc test was used to determine whether specific time points were significantly different from zero. Adjusted p-values (q) were calculated from ANOVA p-values by applying the Benjamini-Hochberg method to correct for multiple comparisons.

### Fuzzy clustering

Differentially-regulated peptides as determined by adjusted ANOVA were clustered using the package Mfuzz (v2.40) in R (v.3.4). The data were standardized so that each peptide had an average value of zero and a standard deviation of one across its time series. The fuzzifier parameter *m* was estimated at 1.75 using the package’s internal mestimate function. The number of clusters *c* was determined by analyzing all *c* from 2 to 20, and identifying no new temporal patterns beyond five clusters. Data were plotted using mfuzz.plot2.

### PANTHER annotations

Three of the five clusters identified using fuzzy c-means clustering had similar temporal patterns and were combined for downstream analysis. Peptides were discretely assigned to one of the three representative clusters based on their highest cluster membership values. Peptides from each cluster were then represented by their respective genes only if all regulated peptides for each gene belonged to a single cluster. These gene-annotated clusters were analyzed for GO term statistical overrepresentation using PANTHER (pantherdb.org) and Reactome (Reactome.org). All annotations with a nominal p-value below 0.05 were included in Supp. Table S2.

### Ingenuity Pathway Analysis

Ingenuity Pathway Analysis (Qiagen) was completed for each individual time point. For each gene, the log_2_ fold change versus zero of the single most changing (absolute value) peptide that passed Dunnett’s post-hoc test was submitted for analysis. For peptides shared by multiple genes, only those with the highest annotation score (Uniprot) were used. The reference set (background) was defined as the entire “User Dataset”, which included all remaining, non-changing genes (with peptides that did not pass Dunnett’s). Only direct relationships were considered and limited to “Experimentally Observed” confidence. All data sources, tissues, cell lines, and mutations were used, but only from the Human species. To indicate EGF redox dynamics over 60 min for the various pathways presented in figure 4, genes were first curated to each pathway to eliminate overlap. Then, the time course of log_2_ fold-changes for peptides belonging to curated genes were added to the pathway heatmap if they were EGF-regulated at one or more timepoint.

### Bioinformatic analysis

The amino acid position in the protein for each cysteine and site-specific annotation (i.e active sites, disulfide type, topology and subcellular localization) was obtained using an in-house python script that searched the UniProtKB flat file (2018_07, 20380 entries). Protein domain and family annotation as well as the start and end position for all human proteins was obtained from PFAM (v31) and used to create a domain specific FASTA database (BLAST v2.2.26). All non-cysteine containing domains and families were excluded and each identified site covered by a domain was annotated. Over-representation was identified by comparing all domains identified in our dataset versus all cysteine-containing domains in the human proteome and validated using a two-sided Fisher’s exact test in R (v.3.3.3) (Supp. Table S4). The combined charge at pH 7 of the amino acids stretch (+/−2) surrounding each cysteine was calculated using the R package Peptides v2.4 (26641634) and compared to the global charge distribution of all cysteines in the identified proteins. Only cysteines that can be theoretically covered by tryptic digestion were considered to prevent a bias for positively charged K/R residues and data analysis were performed using GraphPad Prism (v7.0b) by two-way ANOVA. Relative solvent accessibility (RSA) and secondary structure prediction was performed using NetSurfP-2 (68). An RSA above 25% was considered surface accessible and secondary structures were assigned as helix, turn and beta-strand.

Differentially alkylated peptides were only evaluated when all forms were identified (i.e 2-Ox and both 1-Ox forms). Pearson linear regression was calculated for the 1-Ox peptides across the time course as well the standard deviation at t60.

### Calculating percent cysteine conservation and phylogenetic analysis

The entire Uniprot protein sequence was aligned with Clustal Omega (EMBL) followed by phylogenetic tree generation using Simple Phylogeny (EMBL), both using default parameters. Phylogenetic trees were visualized with iTOL (itol.embl.de) and annotated using domain information from Uniprot and gene family information from the Human Gene Nomenclature Committee.

### Protein structure analysis

PDB structures were visualized with Chimera (UCSF).

### Calculating percent of proteins significantly oxidized in each compartment

Peptides were assigned to gene level as for IPA. Only proteins annotated to a single subcellular location in Uniprot were analyzed since these are sentinels of specific compartments. The percent of significantly oxidized genes was calculated by taking the number of Dunnett positive proteins at each timepoint divided by the total number of sentinels assigned to each organelle.

### Molecular dynamics simulations

Molecular dynamics simulations of GDP-bound KRAS, were run with Gromacs 5.1.1 at 300 K using the AMBER03 force field with explicit TIP3P solvent(1, 16, 40). Simulation parameters for the GDP are described elsewhere (83). Simulations were prepared by placing the starting structure (PDB ID: 4OBE) in a dodecahedron box that extends 1.0 Å beyond the protein in any dimension. The system was then solvated (21490 atoms), and energy minimized with a steepest descents algorithm until the maximum force fell below 100 kJ/mol/nm using a step size of 0.01 nm and a cutoff distance of 1.2 nm for the neighbor list, Coulomb interactions, and van der Waals interactions. For production runs, all bonds were constrained with the LINCS algorithm and virtual sites were used to allow a 4 fs time step (20, 37). Cutoffs of 1.0 nm were used for the neighbor list, Coulomb interactions, and van der Waals interactions. The Verlet cutoff scheme was used for the neighbor list. The stochastic velocity rescaling (*v*-rescale) (9) thermostat was used to hold the temperature at 300 K. Conformations were stored every 20 ps.

The FAST algorithm (102, 103) was used to enhance conformational sampling of states with large pocket openings. Pocket volumes were calculated using the LIGSITE algorithm, as implemented in enspara (70). FAST-pockets simulations were run for 25 rounds, with 10 simulations per round, where each simulation was 40 ns in length (10 μs of aggregate simulation). In addition to ranking based on pocket volumes, a similarity penalty was included to promote conformational diversity in starting structures (104). An MSM was built from the FAST simulation data using enspara (https://github.com/bowman-lab/enspara). The state space was defined using backbone heavy atoms (atoms C, C_α_, C_β_, N, O), which was clustered with a *k*-centers algorithm based on RMSD between conformations until every cluster center had a radius less than 1.2 Å. Following clustering, an MSM was built by row-normalizing the observed transition counts, at a lag-time of 2 ns, with a small pseudo-count as a prior (105). Solvent accessible surface areas for each state in the MSM were calculated using MDTraj and were weighted based on their determined populations (55).

### Phylogenetic analysis of GTPases

A multiple sequence alignment of all human small GTPases was performed with Clustal Omega (https://www.ebi.ac.uk/Tools/msa/clustalo/) and phylogenetic analysis was performed with Simple Phylogeny (https://www.ebi.ac.uk/Tools/phylogeny/simple_phylogeny/). iTOL (https://itol.embl.de) was used to display phylogenetic tree and biochemical properties of each GTPase.

## Acknowledgements

We acknowledge funding and support from R01 CA200893 (J.H), R21 CA138308 (J.H.), R21 CA179452 (J.H.), VR 2015-00656 (S.V.D.P), SSMF P17-0060 (S.V.D.P), the DRC at Washington University (Grant No. 5 P30 DK020579) and AB SCIEX. G.R.B. and M.I.Z. were funded by NIH R01GM124007 and NSF CAREER Award MCB-1552471. A.M. was supported by the T32GM007067-41 T32 training grant.

## Author Contributions

J.M.H. designed the study. J.M.H and M.J.E. carried out the experiments. J.B.B, S.V.D.P, A.D.M, M.I.Z, G.R.B, and J.M.H. performed data processing, analysis and interpretation. J.B.B, S.V.D.P, A.D.M, M.I.Z, G.R.B. and J.M.H wrote the paper.

## List of Abbreviations

DDA: data-dependent acquisition
DIA: data-independent acquisition
EGF: Epidermal growth factor
EGFR: Epidermal growth factor receptor
ER: endoplasmic reticulum
GO: gene ontology
GSH: glutathione
GSSG: glutathione disulfide
H_2_O_2_: hydrogen peroxide
HIF: Hypoxia-inducible factor
IAC: iodoacetamide
IPA: Ingenuity Pathway Analysis (Qiagen)
LMW-PTP: low molecular weight protein tyrosine phosphatase
*m/z*: mass to charge ratio
MS: Mass spectrometry
NADP: Nicotinamide adenine dinucleotide diphosphate
NADPH: Dihydronicotinamide adenine dinucleotide phosphate
NEM: N-ethylmaleimide
OxRAC: Oxidation analysis by resin-assisted capture
PKM: pyruvate kinase
PTPs: protein tyrosine phosphatases
PRDX: peroxyredoxins
pY: pan-phosphotyrosine
ROS: reactive oxygen species
RSA: relative solvent accessibility
RTK: receptor tyrpsine kinase
SDS: sodium dodecyl sulfate
TCEP: tris(2-carboxyethyl)phosphine
UGDH: UDP-glucose 6-dehydrogenase
VCP: valosin-containing protein

---

**Supplemental Figure S1.**
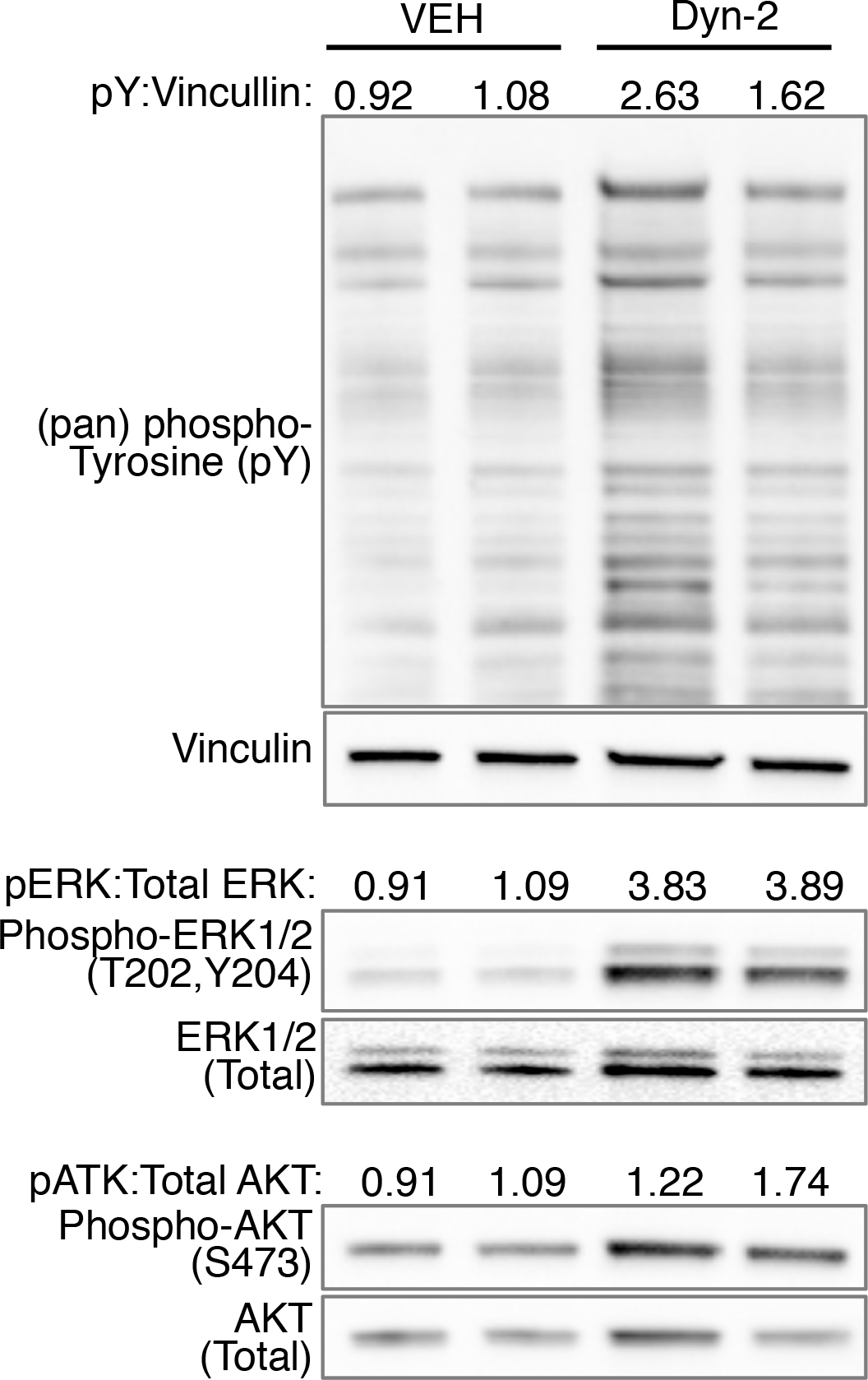
In situ Dyn-2 treatment alters EGFR-dependent signaling. Serum starved A431 cells were prepared as for OxRAC, but the cells were treated with Dyn-2 or vehicle for 1 hour as in (96) (5mM dyn-2 in DMSO, 1 hr). Western blot to pan-phosphotyrosine (pY), vinculin to normalize for protein expression, phospho-T202/Y204 ERK, total ERK, phospho-S473 AKT and total AKT.

**Supplemental Figure S2.**
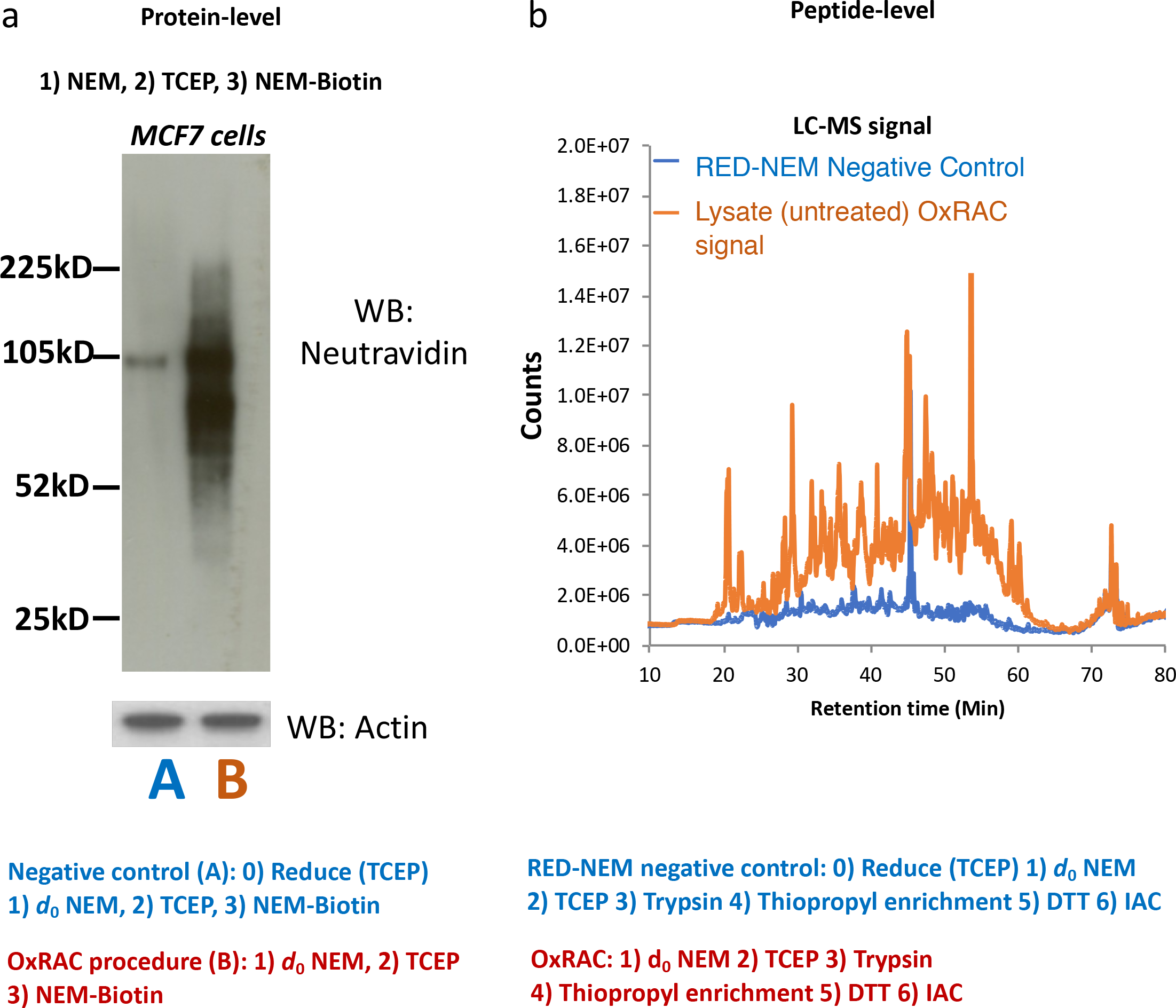
OxRAC method validation and controls. **A)** MCF7 cells were lysed in SDS plus unlabeled NEM as in the OxRAC procedure to block free thiols. After reduction, labeling of previously oxidized cysteines with NEM-biotin generates significant signal when blots are probed with neutravidin-HRP. However, the negative control which is pre-reduced prior to NEM treatment has negligible signal validating complete alkylation during lysis. **B)** Similar experimental design as in A, but performing thiopropyl sepharose enrichment and mass spectrometry. ‘RED-NEM’ samples which are reduced prior to NEM alkylation have very low total ion chromatogram by mass spectromertry and few cysteine peptides identified as compared to samples processed for OxRAC analysis.

**Supplemental Figure S3.**
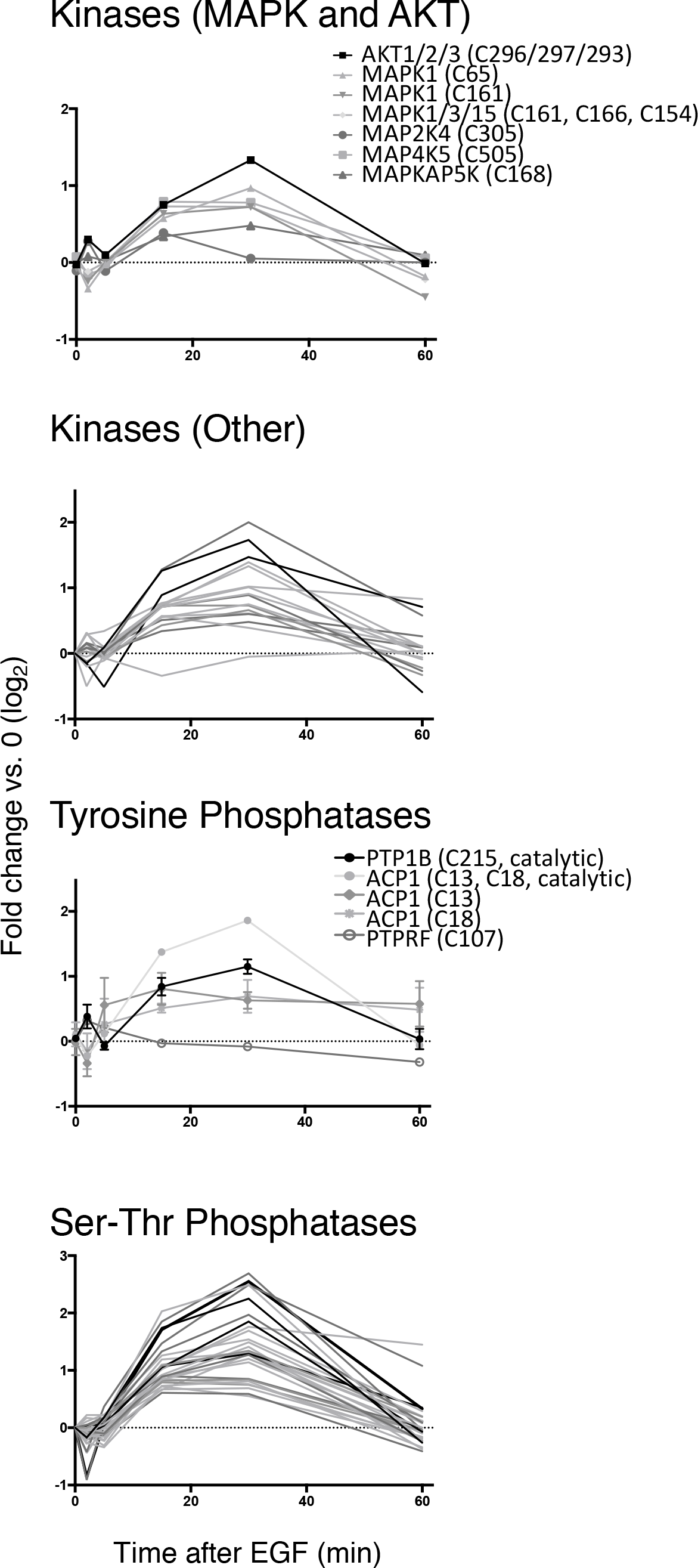
Redox regulation of protein kinases and phosphatases. Time course of redox regulation of cysteines in significantly oxidized peptides assigned to kinases and phosphatases. Each line represents one peptide.

**Supplemental Figure S4.**
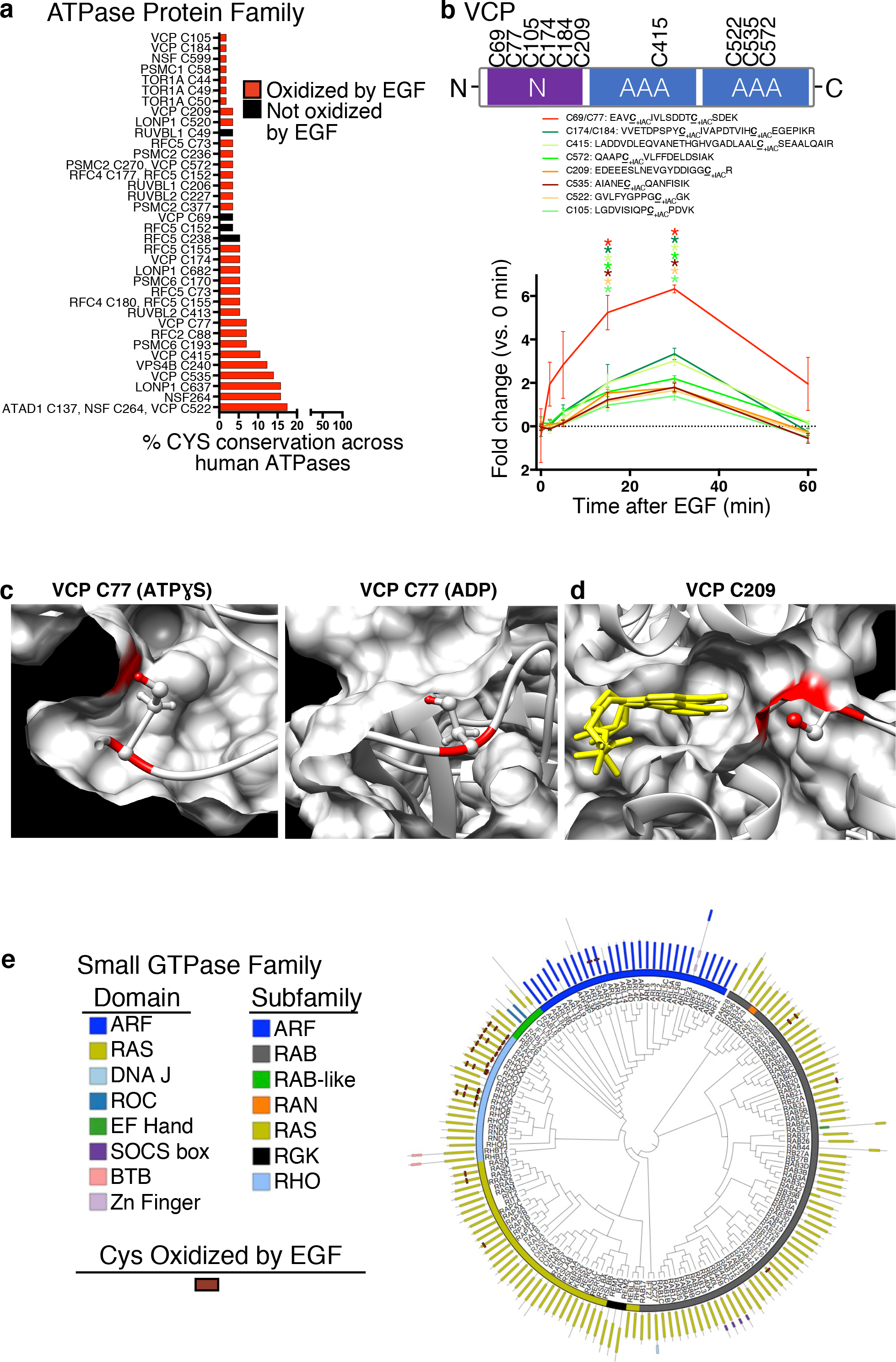
EGF-dependent redox regulation of AAA ATPases and small GTPases. **A)** Amino acid sequence conservation of the cysteines identified in AAA ATPase proteins with significance (q ≤ 0.05) of EGF response indicated. **B)**Time-dependent changes in the oxidation of all cysteines in VCP. Asterisks indicate significance by ANOVA (Dunnett's post hoc test). **C)** Crystal structures of VCP. VCP C77 is solvent accessible in the ATPƔS VCP structure (PDBID: 5FTN) but buried and solvent inaccessible in the ADP bound state (PDBID:5FTL). The proton on the thiol is colored red as is its contribution to the protein surface. **D)** VCP C209 (red) lies at the base of the nucleotide (yellow) binding pocket. **E)** Phylogenic analysis of all human small GTPases with the subfamily (inner ring), domain (outer ring), and location of significantly oxidized cysteines indicated. Line length corresponds to the number of amino acids with the N-terminus facing the center.

**Supplemental Figure S5.**
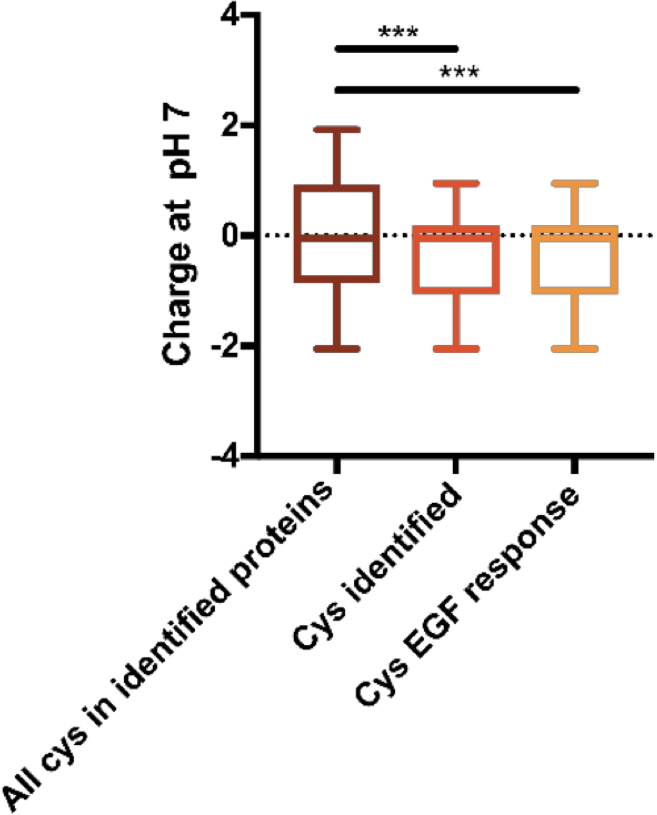
Biochemical properties of the identified oxidized cysteines. Net charge at pH 7 for all cysteines +/−;2 amino acids.

